# Nanoscopy of organelles and tissues with iterative ultrastructure expansion microscopy (iU-ExM)

**DOI:** 10.1101/2022.11.14.516383

**Authors:** Vincent Louvel, Romuald Haase, Olivier Mercey, Marine. H. Laporte, Dominique Soldati-Favre, Virginie Hamel, Paul Guichard

## Abstract

Expansion microscopy is an approach of super-resolution fluorescence microscopy that does not yet achieve the precision of nanoscopy techniques such as single-molecule light microscopy (SMLM). Here, we developed an iterative ultrastructure expansion microscopy approach (iU-ExM), which now matches the SMLM resolution as demonstrated using standard references such as the nuclear pores. Applicable to both cells and tissues, iU-ExM allows a broad research community to access high precision super-resolution microscopy.

## Main Text

In the same way as conventional electron microscopy (EM) at the 20^th^ century^1^, the invention of super-resolution microscopy (SRM), has recently revolutionized our vision of the molecular architecture of the cell^2^. Of all existing methods, single molecule light microscopy (SMLM) approaches, including PALM, STORM, PAINT, MINFLUX, have the highest resolving power with near-nanometer resolution ^3^. However, these methods require advanced equipment and expertise, which limits their use for most laboratories.

The development of expansion microscopy (ExM) in recent years enabled to democratize super-resolution by leveraging the isotropic expansion of the biological samples^4^. This revolutionary methodology based on a 4-fold increase of the distance between molecules, now allows obtaining a resolution in the order of 70 nm with a widefield or confocal microscope ^4^. However, since its original description and despite many protocol variations to further increase the expansion factor ^5^, the effective resolution of ExM alone still does not quite reach that of SMLM approaches.

Here, we undertook to push the resolution of this method by combining two approaches, Ultrastructure Expansion Microscopy (U-ExM)^6^, known to reduce the linkage error and thus increase the labeling precision^7^, with iterative expansion microscopy such as iExM or Pan-ExM methods ^8,9^ in which a first cleavable gel (DHEBA) is used allowing a second round of expansion after re-embedding in a second uncleavable gel (Bis-acrylamide). Initially, we tested whether this first cleavable gel could resist the denaturation step at 95°C as used in the original U-ExM protocol. We found that the DHEBA gel melt at this temperature, probably due to cross-linker cleavage and thus decided to probe its solidity at lower temperatures. We thus tested 85°C that is closer to the 95°C U-ExM denaturation and 73°C, a temperature already validated in the Pan-ExM protocol ^9^. Importantly, at both temperatures, the first cleavable gel remained intact. We next monitored, in the first expanded gel, the expansion achieved under both conditions by measuring the nuclei cross section (**Supplementary Fig. S1a, b**). We found that, as expected when using a higher temperature for the denaturation step, the nuclei were better expanded at 85°C reaching complete expansion as with U-ExM (**Supplementary Fig. S1a, b**). To test whether the molecular expansion was also better preserved at 85°C, we immunolabeled the centriole using both tubulin and NHS-ester ^9,10^ (**Supplementary Fig. S1c**). We found that, while centrioles were not fully expanded at 73°C, they were isotropically expanded at 85°C similarly to what we reported using U-ExM ^6^ (**Supplementary Fig. S1c, d**). Similarly, the conoid, a tubulin fibers based structure of the apicomplexan parasite *Toxoplasma gondii* known to be isotropically expanded using U-ExM ^11^, correctly expanded at 85°C but not at 73°C. Moreover, the fluorescent detection of doublecortin DCX displayed a more defined signal with a higher signal to noise ratio at 85°C (**Supplementary Fig. 1e**). We therefore opted to adapt the U-ExM method using a denaturation step at 85°C that is compatible with the iterative protocol. We next monitored the post-expansion labeling after iterative U-ExM (referred to as iU-ExM thereafter) using either the pan marker NHS-ester or antibodies against tubulin to visualize microtubules. As previously reported, NHS-ester labeled the centriole, despite an incomplete expansion at 73°C compared to 85°C, unveiling its architectural details^9^ (**Supplementary Fig. 1f, g**). In contrast, post-expansion staining of tubulin turned out to be very weak (**Supplementary Fig. 1h, top panels**), a technical issue that could be due to the density of the two gels, which limits the diffusion of antibodies. We therefore tested whether an intermediate staining, i.e. in the first gel, would be retained in the second gel and improve the fluorescent signal. Remarkably, we found that this approach results in a significant intensity increase in the second gel as shown with the fluorescent signal of labeled microtubules (**Supplementary Fig. 1h, bottom panels**). We next further assessed this intermediate staining strategy using *T. gondii* parasites as a reference sample and found that tubulin-based structures in the parasite could be visualized with a full width at half-maximum (FWHM) of 26.7 +/- 2.8 nm, consistent with the reported size of individual microtubules in electron microscopy ^12^ (**Supplementary Fig. 1i**). These results demonstrate that the intermediate staining strategy works similarly to post-expansion labeling with a neglectable linkage error ^7^.

Next, we assessed whether iU-ExM would enable the visualization of the nuclear pore complex (NPC), a stereotypical macromolecular assembly made of two 8-fold symmetrical stacked rings used a reference structure to evaluate SRM performance that remains difficult to resolve using ExM ^13^. We first tested fixation conditions as well as antibody labeling strategies using the validated NUP96-GFP cell line to visualize the NPC (**Fig. 1** and **Supplementary Fig. 2**). Consistent with the previous studies ^13,14^, the best fixation condition to visualize NPC was a pre-extraction-based protocol (see methods) (**Supplementary Fig. 2a-c**). In addition, we found that double antibody labeling using a mix of antibodies directed against NUP96 and GFP gave a better signal and an improved labeling efficiency of NPC showing a greater proportion of 8-fold NPCs (**Fig. 1b, c**; **Supplementary Fig. 2d-i**). Finally, using widefield and confocal microscopes, we demonstrated that iU-ExM not only unveils the 8-fold symmetry of the NPC but also the NUP96 localization in the nuclear and cytoplasmic rings seen from side view, matching the SMLM resolution (**Fig. 1b, c**). Concordantly, the average angle between NUP96 dots is 45.75 +/- 7.8° (**Fig. 1d**), further supporting the 8-fold distribution of the NUP96 as described in SMLM methods ^13,15,16^.

**Figure 1.**
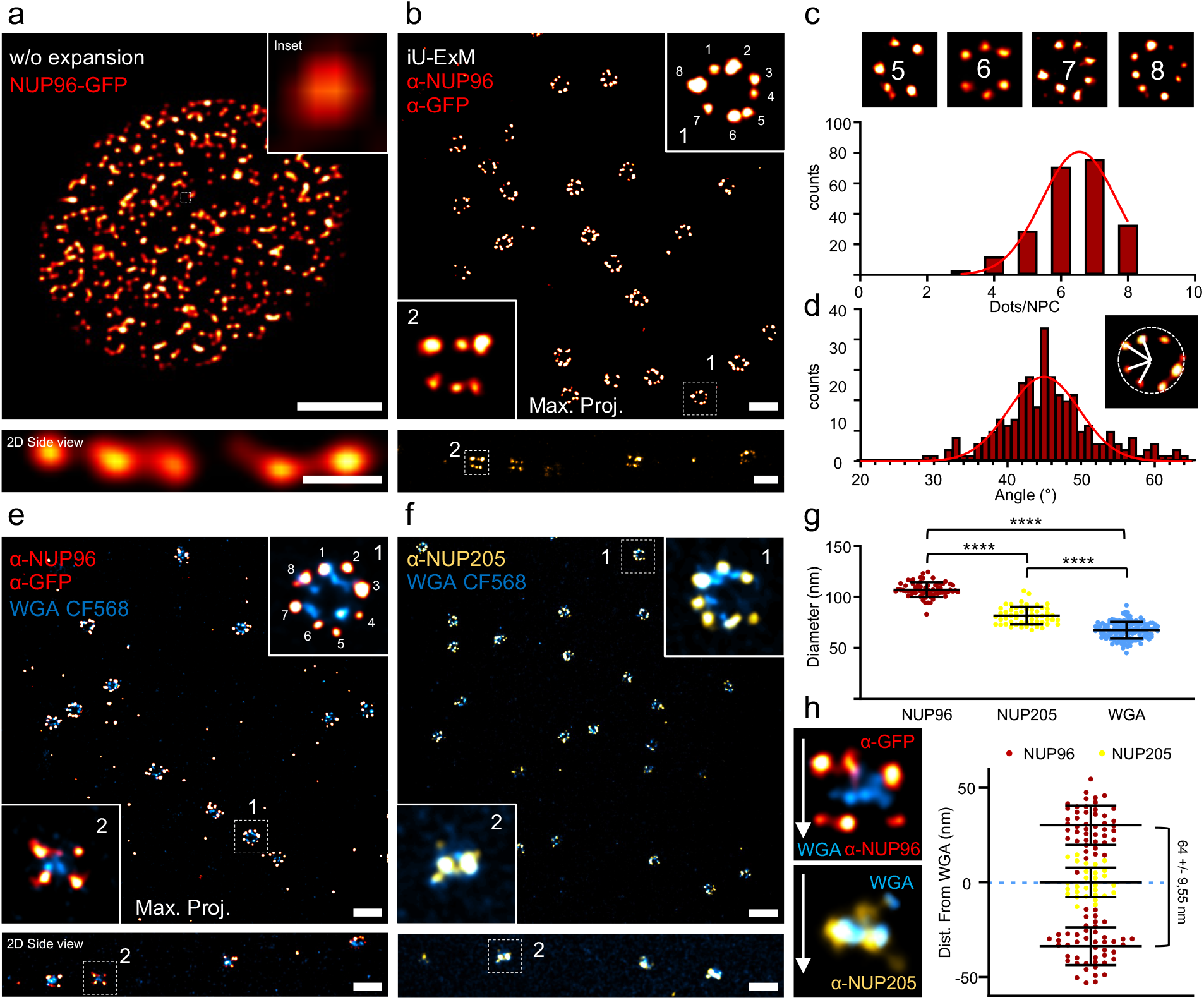
iU-ExM reveals the 8-fold organization of the human Nuclear Pore Complexes. **(a)** Confocal image of NUP96-GFP positive nucleus illustrating the endogeneous GFP signal (red hot) without expansion. Scale bar: 4 µm (upper image), 500 nm (lower). (**b**) iU-ExM confocal image of NPC labelled with α-GFP and α-NUP96 primary antibody mix, revealing the NUP96 8-fold pattern (inset 1) as well as the cytoplasmic and nucleoplasmic outer rings of the NPC (inset 2). Scale bars: 200 nm. (**c**) Quantification of the number of dots per NPCs. N= 224 NPCs from 3 independent experiments. Red continuous line: non-linear gaussian regression (R^2^= 0.99). (**d**) Histogram of the angles between neighboring NUP96 dots in iU-ExM imaged NPCs. N= 191 NPC from 3 independent experiments (average +/- standard error = 45.75 +/- 7.8°). Red continuous line: non-linear gaussian regression (R^2^= 0.86, Mean +/- side deviation = 44.96 +/- 4.8°). (**e, f**) Dual color iU-ExM confocal images of nuclei stained for WGA (NPC inner ring, cyan) together with NUP96 (red hot, e) or NUP205 (yellow, f). Scale bars: 240 nm. (**g**) Quantification of the diameter of NUP205, NUP96 and WGA in top viewed NPC. Note that the expanded NUP96 diameter was divided by 107 nm, which represents the published NUP96 value ^13^ to determine the expansion factor (Exp. F= 16.8X). This value was next applied to the expanded measurements of NUP205 and WGA. NUP96: N= 66 (average +/- standard error = 107 +/- 7.2 nm), NUP205: N= 54 (average +/- standard error = 81.6 +/- 8.7 nm), WGA: N=138 (average +/- standard error = 67.3 +/- 8.2 nm), from 3 independent experiments. P-values: ****: p<0,0001. One-way ANOVA. (**h**) Quantification of the NUP96 and NUP205 position relative to the WGA signal seen in side view NPC. NUP96: N= 46 (average +/- standard error = 64 +/- 9.5 nm), NUP205: N=46 (average +/- standard error = 0.5 +/- 7.6 nm) from 3 independent experiments.

We next tested dual staining that often remains challenging in SMLM ^5^. To do so, we stained expanded human U2OS cells with either NUP96 or NUP205, an inner NPC component and WGA, a marker of the nuclear pore inner ring (**Fig. 1e-h**). We found that iU-ExM enables to discriminate between the inner and outer parts of the NPC, with NUP96 located on the nuclear and cytoplasmic outer rings, while NUP205 and WGA decorated the inner part of NPC (**Fig. 1e-h**). We next calibrated the expansion factor achieved by iU-ExM using the previously reported NUP96 diameter of 107 nm in SMLM and cryo-electron microscopy (cryo-EM) ^17^, resulting in a ±17-fold expansion. With this calibrated expansion factor, the NUP96 cytoplasmic and nucleoplasmic rings were 64 nm +/- 9.55° apart (**Fig. 1h**). To further improve the imaging of NPCs, we turned to isolated nuclei using the NUP96-GFP cell line (**Supplementary Fig. 3**). We found that iU-ExM similarly enabled revealing the canonical features of NPC, 8-fold symmetry and a measured NUP96 thickness of 52 nm +/- 7.98° (**Supplementary Fig. 3a-g and Movies 1 and 2**), coherant with the cryo-ExM structure and SMLM measurements ^13,17^.

To further investigate the potential of iU-ExM to reveal the organization of cellular assemblies, we next focused on tubulin-based structures, the *Chlamydomonas reinhardtii* basal body (BB) and flagella, as well as the conoid of *T. gondii*. These structures can serve as molecular rulers as they have canonical dimensions of, 225 nm wide and 500 nm long for the BB ^18^, and 280 nm in length and 380 nm in diameter for the spiraling tubulin fibers of the conoid ^19^. iU-ExM applied on isolated BB stained for αβ-tubulin and imaged with a widefield microscope unambiguously revealed their 9-fold microtubule triplet/doublet chirality with a correct length/diameter ratio as observed previously ^6^ (**Supplementary Fig. 4a-c**). Furthermore, whole *C*.*reinhardtii* cell expansion using iU-ExM enabled to resolve the microtubule doublets and the central pair that compose the axoneme *in vivo* (**Supplementary Fig. 4d**). These results demonstrate that iU-ExM alone reaches a resolution similar to the previously published combination of U-ExM combined to STED ^6^. Next, we expanded *T. gondii* parasites (**Fig. 2** and **Supplementary Fig. 5**) and found that iU-ExM can reveal the spiral structure of the 40 nm apart conoid tubulin fibers (**Fig. 2b**), a structural feature which to date has only been observed by electron microscopy^20^ (**Fig. 2c**). Moreover, the 9-fold symmetry of *T. gondii* centrioles (**Supplementary Fig. 5a-c**) and the 22 cortical microtubules branching from the apical polar ring (APR) ^19^ now became clearly resolvable (**Fig. 2b, d**). With this nanometric resolution, we undertook to probe the potential for a molecular mapping of the conoid using iU-ExM. To do so, we monitored the spatial distribution of either SAS6-L or DCX relative to tubulin, two proteins previously identified as components of the conoid ^21,22^ (**Fig. 2e-i** and **Supplementary Fig. 5d, e**). We uncovered that SAS6-L is located in between the tubulin fibers, 4.5 nm towards the conoid lumen relative to the fibers, while DCX is located 15 nm from the outer surface of the conoid fibers (**Fig. 2e-i** and **Supplementary Fig. 5d, e**), demonstrating the resolution power of our approach. In addition, we tested whether iU-ExM could be compatible with the 10-fold TREx expansion protocol ^23^ to further increase the expansion factor of the second gel (**Supplementary Fig. 6**). We found that both the conoid and *Chlamydomonas* isolated BB could be expanded around 25-fold, allowing to further increase the apparent structural resolution obtained using a widefield microscope.

**Figure 2.**
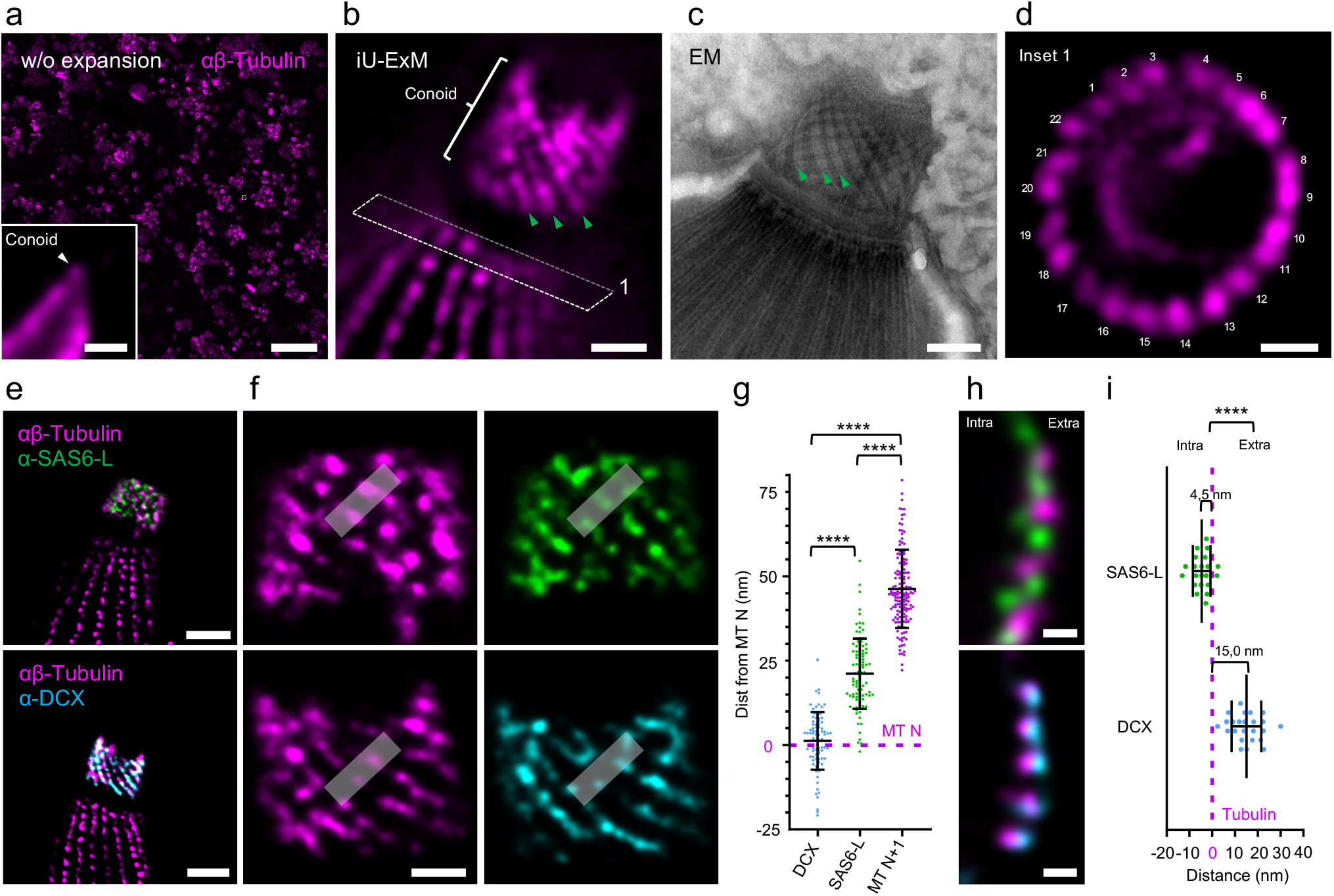
Molecular organization of the conoid in *Toxoplasma gondii* tachyzoites. (**a**) Full field of view widefield image of non-expanded *T. gondii* tachyzoites labelled with tubulin antibodies (magenta). Inset: one parasite with the conoid region (white arrowhead). Scale bars: 30 µm (full picture), 1 µm (inset). (**b**) iU-ExM widefield image unveiling the spiral shape of the tubulin fibers of the conoid (green arrowheads). Scale bar: 100 nm. (**c**) Electron microscopy image of a conoid with the tubulin fibers visible (green arrowheads). Scale bar: 100 nm. (**d**) iU-ExM widefield image of the region marked by (1) in panel b revealing the 22 cortical microtubules branching the apical polar ring (APR). Scale bar: 100 nm. (**e**) iU-ExM confocal images of the conoid stained for either SAS6-L (green, upper images) or DCX (cyan, bottom images) and tubulin (magenta). Scales bars: 200 nm. (**f**) Zoom in of the conoid (from dashed square in panel e) highlighting the distribution of SAS6-L (green) and DCX (cyan) relative to tubulin (magenta). Scale bars: 80 nm. (**g**) Quantification of the position of SAS6-L and DCX relative to the tubulin fibers. DCX: N= 80 (average +/- standard error = 1.2 +/- 8.5 nm), SAS6-L: N= 98 (average +/- standard error = 21.1 +/- 10.4 nm), MT N+1: N = 162 (average +/- standard error = 46.3 +/- 11.6 nm) from 3 independent experiments. (**h**) Side view of the conoid stained for SAS6-L (green) or DCX (cyan) and tubulin (magenta). Scale bars: 40 nm. (**i**) Quantification of the position of SAS6-L and DCX relative to the tubulin fibers. DCX: N= 24 (average +/- standard error = 15.0 +/- 6.5 nm), SAS6-L: N= 21 (average +/- standard error = -4.6 +/- 3.9 nm) from 3 independent experiments. Note that the expansion factor was calculated based on the expanded conoid diameter in the apical part divided by 380 nm^19^. P-values: ****: p<0,0001. Non parametric Kruskal Wallis test. (g) and student t-test (i).

Finally, we wanted to assess the general applicability of iU-ExM and monitored the expansion of membrane-based organelles such as the mitochondria and the endoplasmic reticulum (ER) (**Fig. 3a-d** and **Supplementary Fig. 7**). We found that iU-ExM expanded and preserved PFA-GA fixed mitochondria stained with NHS-ester ^9^, revealing the correct cristae spacing of 85.10 nm +/- 14.34 ^24^ (**Fig. 3a-c**). To visualize the ER, cryo-ExM, which best preserves cellular structures in their native state ^25^, was coupled to iU-ExM (**Fig. 3d** and **Supplementary Fig. 7**). Importantly, we could image the ER with utmost precision, revealing the hollow ER tubules at 45 nm using a confocal microscope **(Fig. 3d, e)** as only previously seen using SMLM ^26^. Lastly, we undertook to test whether iU-ExM was also compatible with tissue expansion. To test this, we expanded mouse retinal tissue stained for tubulin and the centriolar protein POC5, that we recently showed underlines the inner wall of the connecting cilium (CC) of photoreceptor cells, using U-ExM ^27^ (**Fig. 3f-j** and **Supplementary Fig. 8**). We found that iU-ExM works perfectly in tissue and further increase the resolution obtained using U-ExM, revealing the 9-fold symmetrical organization of POC5 at the CC but also at the underlying basal body (**Fig. 3i, j** and **Supplementary Fig. 8c, d**). Altogether, this demonstrates that iU-ExM enables nanoscale imaging resolution from isolated macromolecules to tissue.

**Figure 3.**
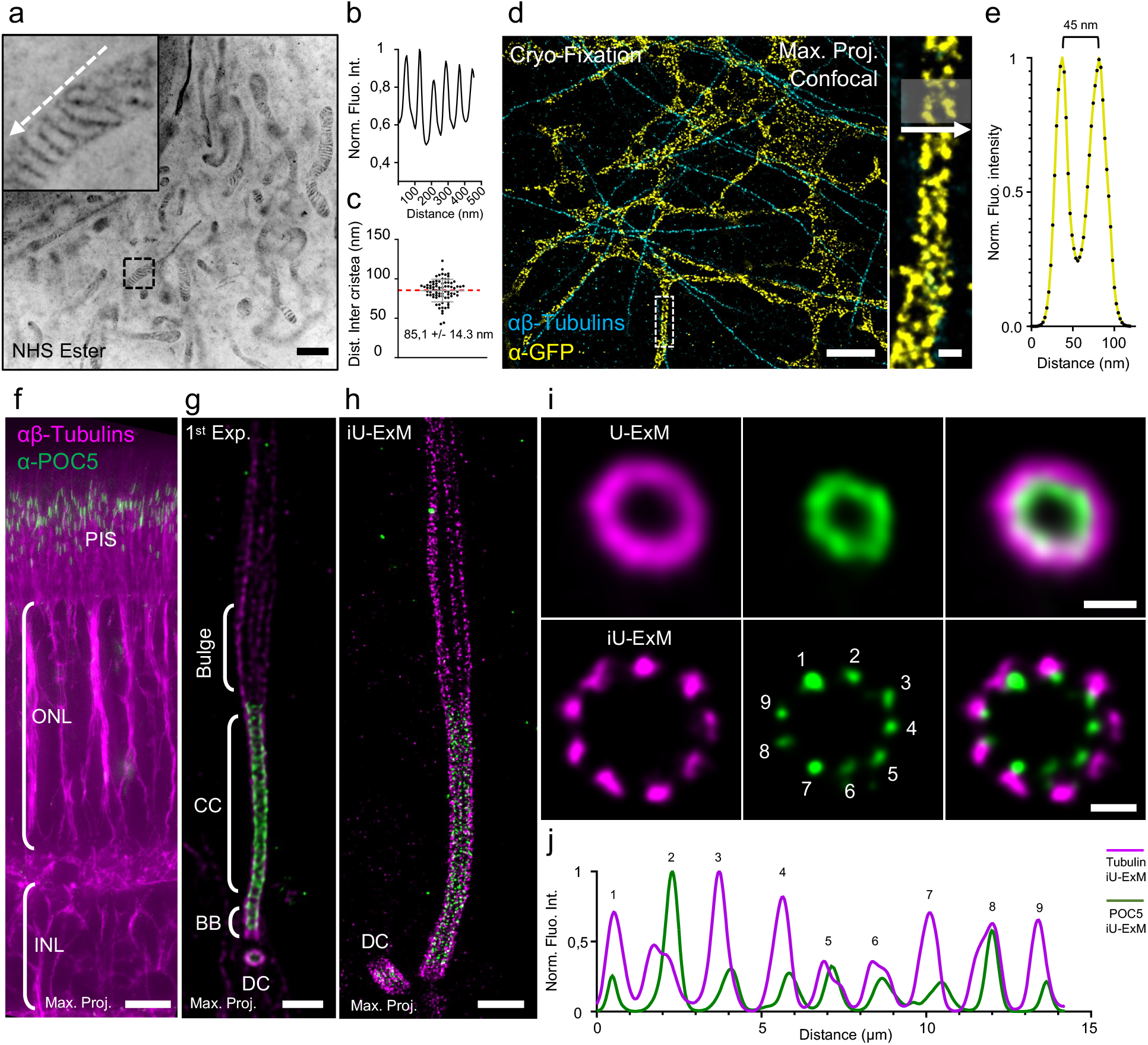
iU-ExM is compatible with tissue expansion and membraneous organelles. (**a**) iU-ExM widefield image of a U2OS cell fixed with 3% PFA, 0.1% GA and stained with NHS-ester to label the mitochondria. Note that iU-ExM allows the visualization of individual cristae (inset). Scale bar: 1 µm. (**b**) Plot profile corresponding to the white dashed arrow in a. (**c**) Quantification of the inter-cristae distance, N= 78 (average +/- standard error = 85.1 +/- 14.3 nm) from 2 independent experiments. (**d**) iU-ExM confocal image of cryo-fixed U2OS cells expressing Sec61β-GFP and labelled with tubulin and GFP, showing the double layer of an endoplasmic reticulum tubule. Scale bar: 500 nm (full image), 50 nm (inset). (**e**) Plot profile of the grey area in the inset. (**f**) widefield image with a 20X objective of an iU-ExM mouse retina tissue stained for tubulin (magenta) and POC5 (green), highlighting the preservation of the different retinal layers. PIS=Photoreceptor Inner Segment, ONL= Outer Nuclear Layer, INL= Inner Nuclear Layer. Scale bar: 25 µm. (**g**) Confocal image of a photoreceptor connecting cilium labelled with tubulin (magenta) and POC5 (green) after the first expansion. CC: Connecting Cilium, BB: Basal Body, DC: daughter centriole. Scale bar: 400 nm. (**h**) iU-ExM confocal image of a photoreceptor stained for tubulin (magenta) and POC5 (green). Scale bar: 400 nm. (**i**) Comparison of top view confocal images of the CC expanded using U-ExM (left) or iU-ExM (right). Note the 9-fold organization of the microtubule doublets as well as the POC5 signal revealed undoudtedly after iU-ExM. Scale bars: 100 nm (U-ExM), 50 nm (iU-ExM). (**i**) Polar transform plot profile for POC5 (green) and tubulin (magenta) signals from the iU-ExM (continuous) top view images of the CC.

## Acknowledgments

We thank Benita Wolf for critical comments on the manuscript. We thank the Kostic and Arsenijevic laboratories (Hospital Jules Gonin, Lausanne, Switzerland) for providing the mouse retinas used in this study (authorization VD1367). We also thank Yanis Bryois for his technical support. This work is supported by the ERC StG 715289 (ACCENT), the Swiss National Foundation (SNSF) 310030_205087 attributed to P.G. and V.H. and 310030_185325 attributed the DSF, as well as the Pro Visu Foundation attributed to P.G andV.H.

## Author Contributions

V.L. performed all the experiments described in the paper, with the help of R.H. for the expansion of *T. gondii* strains, O.M. with retina, and M.H.L. for the cryo-expansion microscopy. R. H. and D.S-F. generated the SAS6-L and DCX strains and provided all the *T. gondii* strains used in this study. The EM image shown in Fig. 2. was produced by Dr. Bohumil Maco. V.H. and P.G. conceived, designed and supervised the project. P.G. and V.H. wrote and revised the final manuscript with inputs from all authors.

## Competing financial interests

Authors declare no competing interests.

## Figure Legends

**Supplementary Figure 1.**
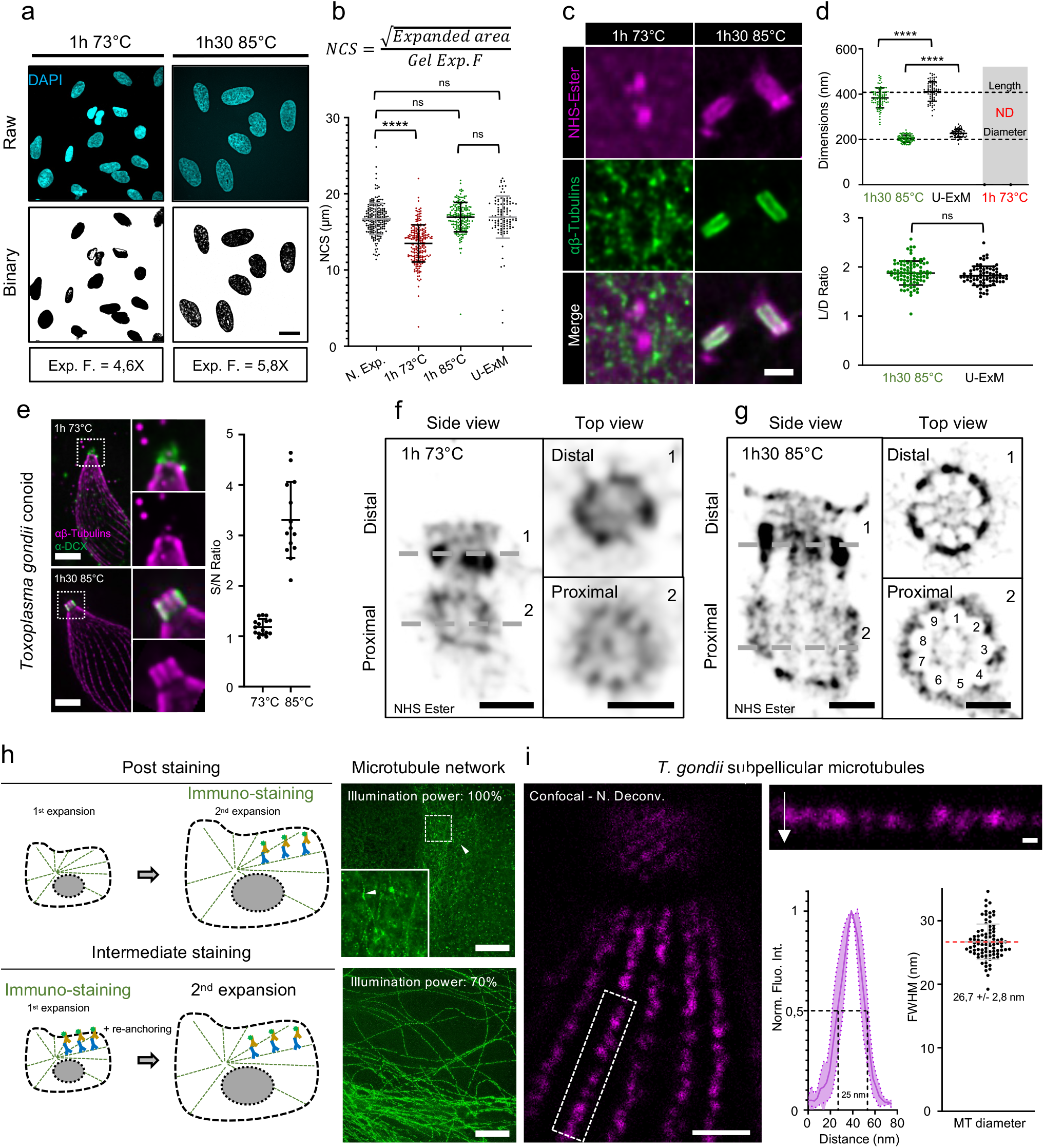
iU-ExM development. **(a)** Representative images of single expanded nuclei from U2OS unfixed cells stained for DAPI in the indicated denaturation conditions. Measurements were performed using the Binary and Area measurement tools of Fiji^28^. Scale bar: 100 μm expanded. **(b)** Quantification of the Nuclei Cross Section (NCS) at different denaturation conditions in unfixed U2OS. NCS was calculated by dividing the square root of the nuclei area with the gel expansion factor measured by the size of the gel before and after expansion (see methods). A lower NCS than the non-expanded value indicates a non-complete nuclear expansion. N. Expanded: N= 215 (average +/- standard error =16.7 +/- 2 μm), 1h 73°C: N= 227 (average +/- standard error = 13.5 +/- 2.4 μm), 1h30 85°C: N= 189 (average +/- standard error = 16.9 +/- 1.9 μm), U-ExM: N= 110 (average +/- standard error = 16.9 +/- 2.8 μm) from 3 independent experiments. (**c**) Representative widefield images of centrioles stained for tubulin (green) and NHS-ester (magenta) after the first expansion (unfixed U2OS). Scale bar: 300 nm. (**d**) upper graph: Quantification of the centriole length and diameter in the indicated conditions corrected by the expansion factor of the gel in unfixed U2OS cells. 1h30 85°C: N= 87 centrioles (average length +/- standard error = 383.9 +/- 43.9 nm, average diameter +/- standard error = 204 +/- 15.5 nm), 1h30 95°C (U-ExM): N= 73 centrioles (length: average +/- standard error = 411.7 +/- 43 nm, diameter: average +/- standard error = 226.7 +/- 17.3 nm) from 3 independent experiments. Note that measurements for the 73°C condition could not be done as the centrioles were not fully expanded nor sufficiently stained. Lower graph: Length/Diameter ratio between the two denaturation conditions highlighting isotropic expansion. 1h30 85°C: N= 87 (average +/- standard error = 1.9 +/- 0.2), 1h30 95°C (U-ExM): N= 73 (average +/- standard error = 1.8 +/- 0.2) from 3 independent experiments. (**e**) Representative confocal images of an expanded (first gel, unfixed) *T. gondii* tachyzoite conoid stained for tubulin (magenta) and doublecortin DCX (green) after 1h 73°C (upper panel) or 1h 85°C (lower panel), maximum projection. Note that the denaturation temperature impacts the full expansion of the conoid structure (insets). Scale bar: 400 nm. Graph: Signal-to-noise ratio in the conoid between the 2 denaturation conditions. 73°C: N= 15 (average +/- side deviation= 1.2 +/- 0.2), 85°C: N= 14 (average +/- side deviation= 3.3 +/- 0.75) from one experiment. (**f, g**) Representative confocal images of iU-ExM U2OS centrioles stained for NHS-ester in the indicated denaturation conditions (f, 73°C) and (g, 85°C) in unfixed conditions. Left: side view, right: top view. Note the 9-fold symmetry visible in both proximal and distal regions of the centriole. We hypothesize that NHS-ester does not label the microtubule triplets and that the footprint of the microtubule wall is visible. Scale bars: 100 nm. Note the difference of resolution with iU-ExM, and the difference in size relatively to the scale bar, underlying the incomplete expansion at 73°C highlighted in (b). (**h**) Scheme of post- and intermediate-staining strategies and representative widefield images of PFA/GA fixed expanded U2OS cells. Scale bars: 500 nm. (**i**). Linkage error evaluation on MT width showing neglectable impact of the linkage error supporting the hypothesis that the 2^nd^ expansion does not increase the linkage error. Scale bar: 80 nm (left image), 25 nm (inset). MT diameter: N=86 (average +/- standard error =26.7 +/- 2.8 nm) from 3 independent experiments. P-value: ****: p<0,0001. One way ANOVA (b,d), student t-test (d, lower graph).

**Supplementary figure 2.**
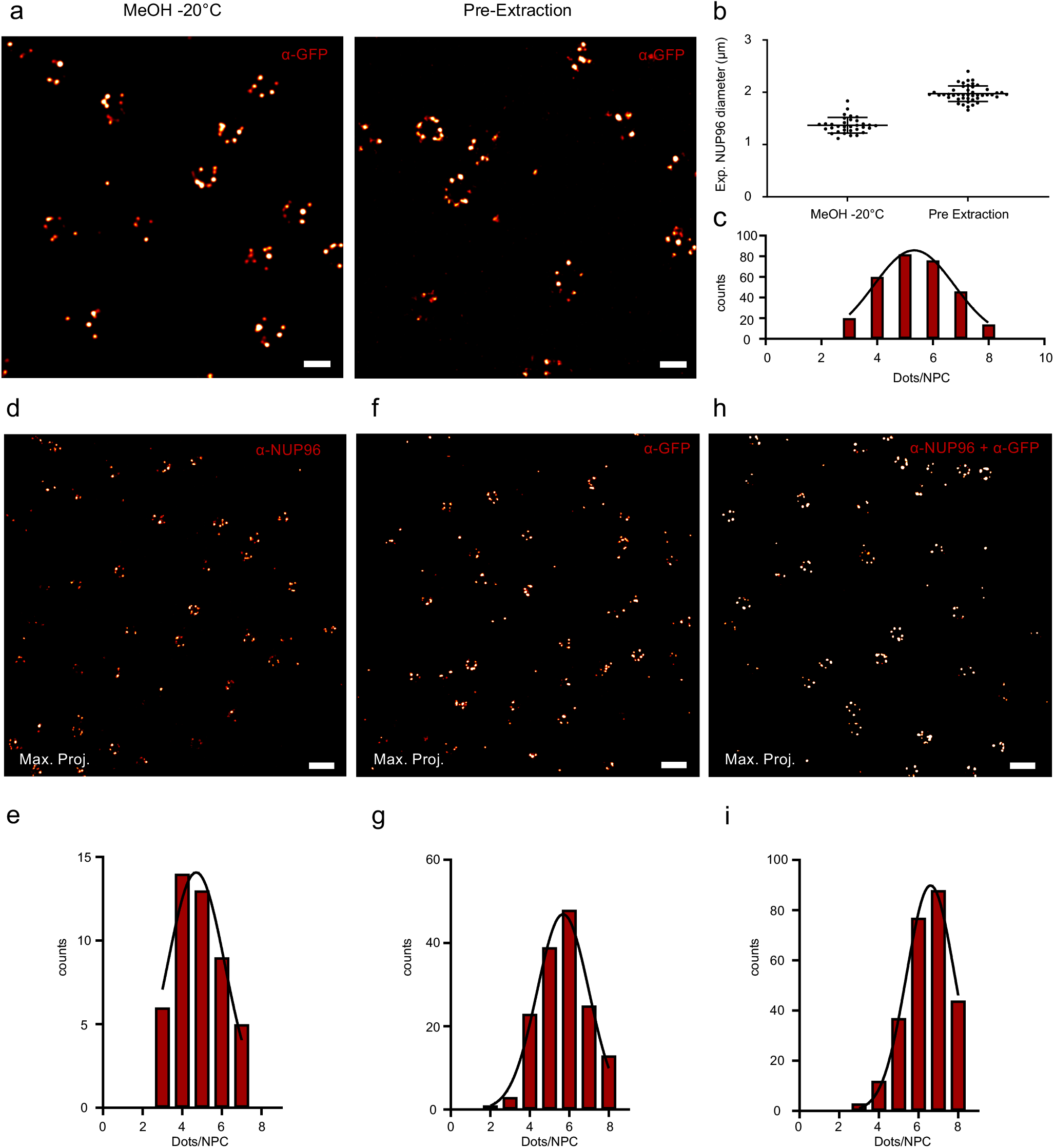
Optimizations of nuclear pores fixations and staining. (**a**) iU-ExM confocal images of U2OS NUP96-GFP cells fixed with Methanol -20°C (MeOH, left) or the pre-extraction protocol from ^13^ (right) and stained with α-GFP antibody. Scales bars: 2 μm non corrected with expansion factor (left right). (**b**) Quantification of the expanded diameter of NUP96-GFP cells fixed with methanol fixation or after pre-extraction. Pre-extraction: N= 45 (average +/- standard error = 1.9 +/- 0.15 μm), MeOH: N=33 (average +/- standard error = 1.4 +/- 0.15 μm) from 3 independent experiments. (**c**) Counts of the number of NUP96 spots per NPC under MeOH fixation, N= 149 NPCs from 3 independent experiments. Continuous black line: gaussian regression (R^2^= 0.99). (**d, f, h)** iU-ExM representative confocal images of pre-extracted U2OS NUP96-GFP cells stained with either primary antibody, α-NUP96 (d), α-GFP (f), or a combination of both antibodies (h) Scales bars: 240 nm. (**e, g, i)** Quantification of the number of NUP96 dots per NPC with either α-NUP96 antibody (e), α-GFP antibody (g), or a combination of both antibodies (i). α-NUP96: N= 47 NPCs from one experiment. Continuous black line: gaussian regression (R^2^= 0.92). α-GFP: N= 152 NPCs from 2 independent experiments. Continuous black line: gaussian regression (R^2^= 0.98). MIX: N= 263 NPCs from 3 independent experiments (Note that this graph is identical to Fig. 1d and serves solely for comparison purposes). Continuous black line: gaussian regression (R^2^=0.99).

**Supplementary Figure 3.**
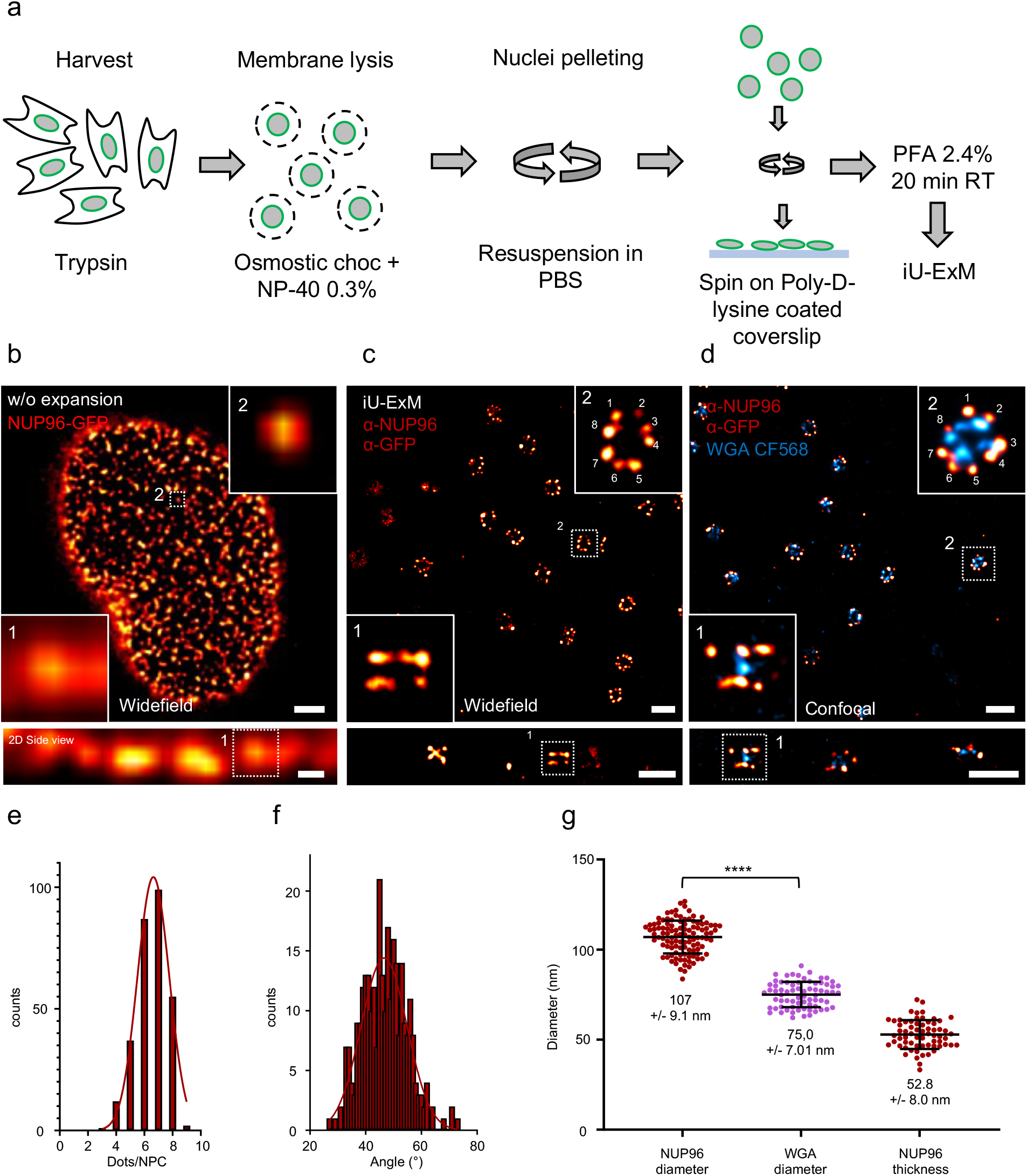
iU-ExM on isolated nuclei. **(a)** Scheme of the nuclei isolation protocol from ^29^. (**b)** Widefield image of an isolated nuclei from NUP96-GFP cells (GFP signal, red hot) (63x, 1.4 NA objective). Scale bars: 2 µm (upper image), 500 nm (lower image, side view). **(c)** iU-ExM widefield image of NPCs stained for NUP96 (red-hot, antibody mix anti-GFP and anti-NUP96) (100x, 1,4 NA objective). Scale bars: 200 nm. **(d)** iU-ExM dual color confocal image of NPC stained with WGA-CF568 (cyan) and NUP96 (red hot, antibody mix). Scales bars: 200 nm. **(e)** Estimation of labelling efficiency by counting the number of NUP96 spots per NPC. N= 293 NPC from 3 independent experiments. Red continuous line: gaussian regression (R^2^= 0.98). **(f)** Quantification of the angle between neighboring NUP96 dots in NPCs. N= 292 NPCs (average +/- standard error = 46.6 +/- 7.9°) from 3 independent experiments. Red continuous line: gaussian regression (R^2^= 0.82, Mean +/- side deviation= 46.6 +/- 8.2 °). (**g)** Quantification of the diameters of the WGA and NUP96 signals and of the distance between the nucleoplasmic and cytoplasmic NUP96 rings (NUP96 thickness). Note that the reported NUP96 diameter of 107 nm is used as a proxy to fix the expansion factor. NUP96 diameter: N= 108 (average +/- standard error = 107 +/- 9.1 nm), WGA: N= 71 (Average +/- standard error = 75 +/- 7 nm), NUP96 thickness: N= 65 (Average= = 52 +/- 7.9 nm), from 3 independent experiments. ****: P-value < 0,0001, Mann-Whitney test.

**Supplementary figure 4.**
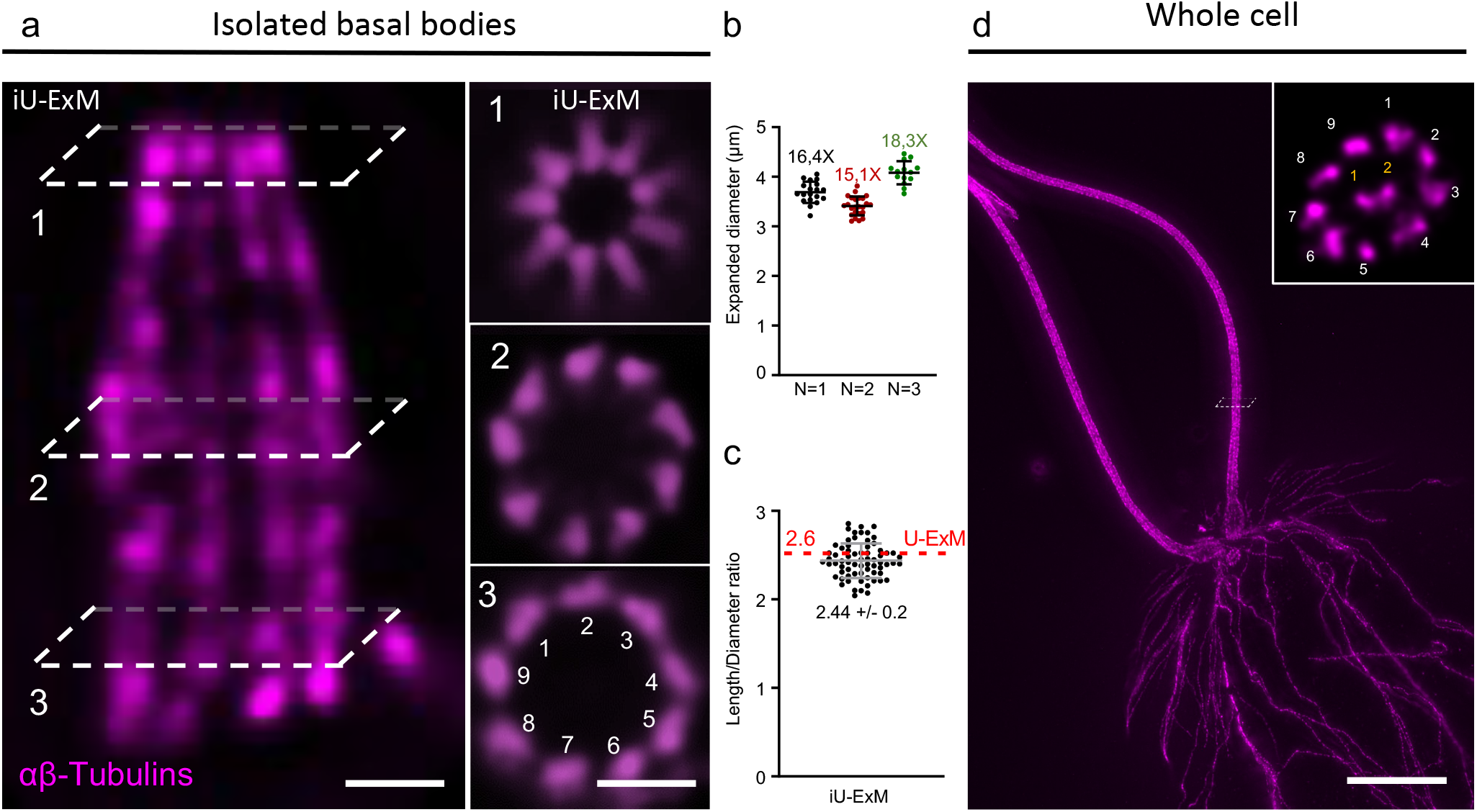
iU-ExM on isolated *Chlamydomonas reinhardtii* centrioles. (**a)** Representative widefield image of an isolated basal body (BB) from *C. reinhardtii* expanded with iU-ExM and stained with tubulin (magenta). Left image, BB from side view with microtubules blades seen longitudinally. Right images: BB top views at the indicated z-position from the dashed boxes on the left image. Scales bars: 100 nm. (**b)** Quantification of the diameter of the proximal area of the basal body for 3 independent experiments. The average value is divided by 225 nm (centriole diameter^30^) to obtain the expansion factor. N= 1 (N= 19 centrioles, average +/- standard error= 3.7 +/- 0.2 μm), N= 2 (N= 24 centrioles, Average +/- standard error= 3.4 +/- 0.2 μm), N= 3 (N= 14 centrioles, average +/- standard error= 4.1 +/- 0.2 μm). (**c)** Length over diameter ratio, the red line shows the value obtained in U-ExM ^6^. N= 69 (Average +/- standard error = 2.4 +/- 0.2) centrioles from 3 independent experiments. (**d)** Widefield image of *C. reinhardtii* cell labelled with anti-tubulin antibodies (magenta). Top inset: (from the white dashed box) top view of the flagella revealing the canonical 9+2 pattern. Scale bar: 1 µm.

**Supplementary figure 5.**
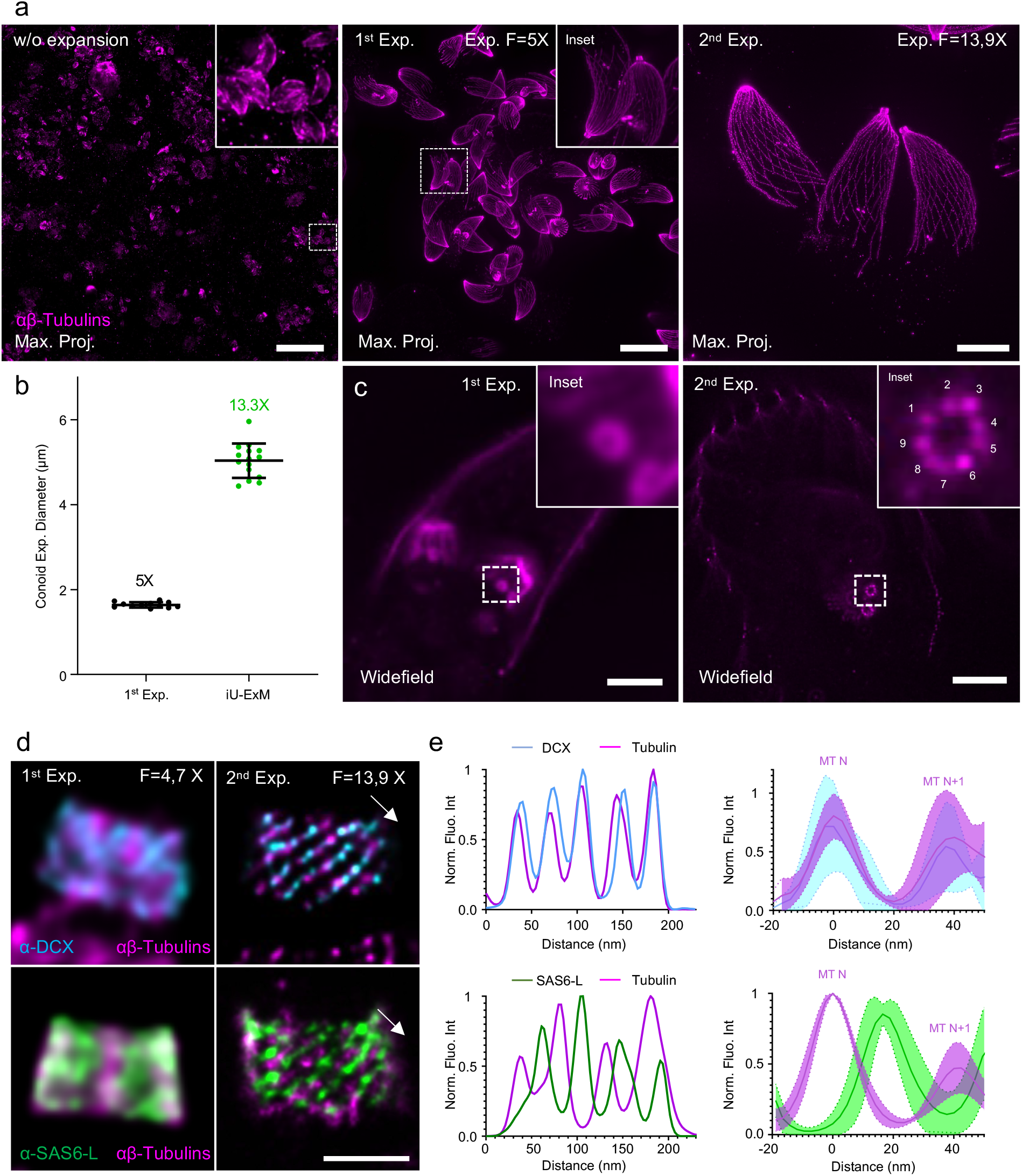
iU-ExM reveals the *Toxoplasma gondii* conoid and centriole ultrastructures. **(a)** Widefield image (full field of view) of non-expanded (left), expanded once (middle) or iU-ExM expanded (right) *T. gondii* tachyzoites stained for tubulin (magenta) (63x 1,4 NA objective). Scale bars: 30 µm (left), 6 µm (middle), 2 µm (right). (**b)** Quantification of the conoid diameter at half its length in the indicated conditions. The expansion factor is calculated by dividing the average diameter value by 380 nm. 1^st^ Expansion (1^st^ Exp.): N= 12 conoids (average +/- standard error= 1.6 +/- 0.1 μm), iU-ExM: N= 15 conoids (average +/- standard error= 5.0 +/- 0.4 μm), from the same experiment. (**c)** Representative widefield images of single (left) or iU-ExM (right) expanded *T. gondii* revealing the 9-fold organization of the centriolar microtubule wall. Scale bars: 4 µm (left), 1.5 µm (right). (**d)** Confocal images of single (left) or iU-ExM (right) expanded *T. gondii* conoids stained for DCX (cyan) or SAS6-L (green) and tubulin (magenta). Scale bar: 200 nm. (**e)** Left: plot profiles in the diagonal (white arrow in d) of the conoid expanded with iU-ExM showing the position of DCX and SAS6-L relative to the tubulin fibers. Right: Pool of 6 plot profiles from different images to illustrate the variability. MT N: tubulin fibers at one given position; MT N+1: neighboring tubulin fibers.

**Supplementary figure 6.**
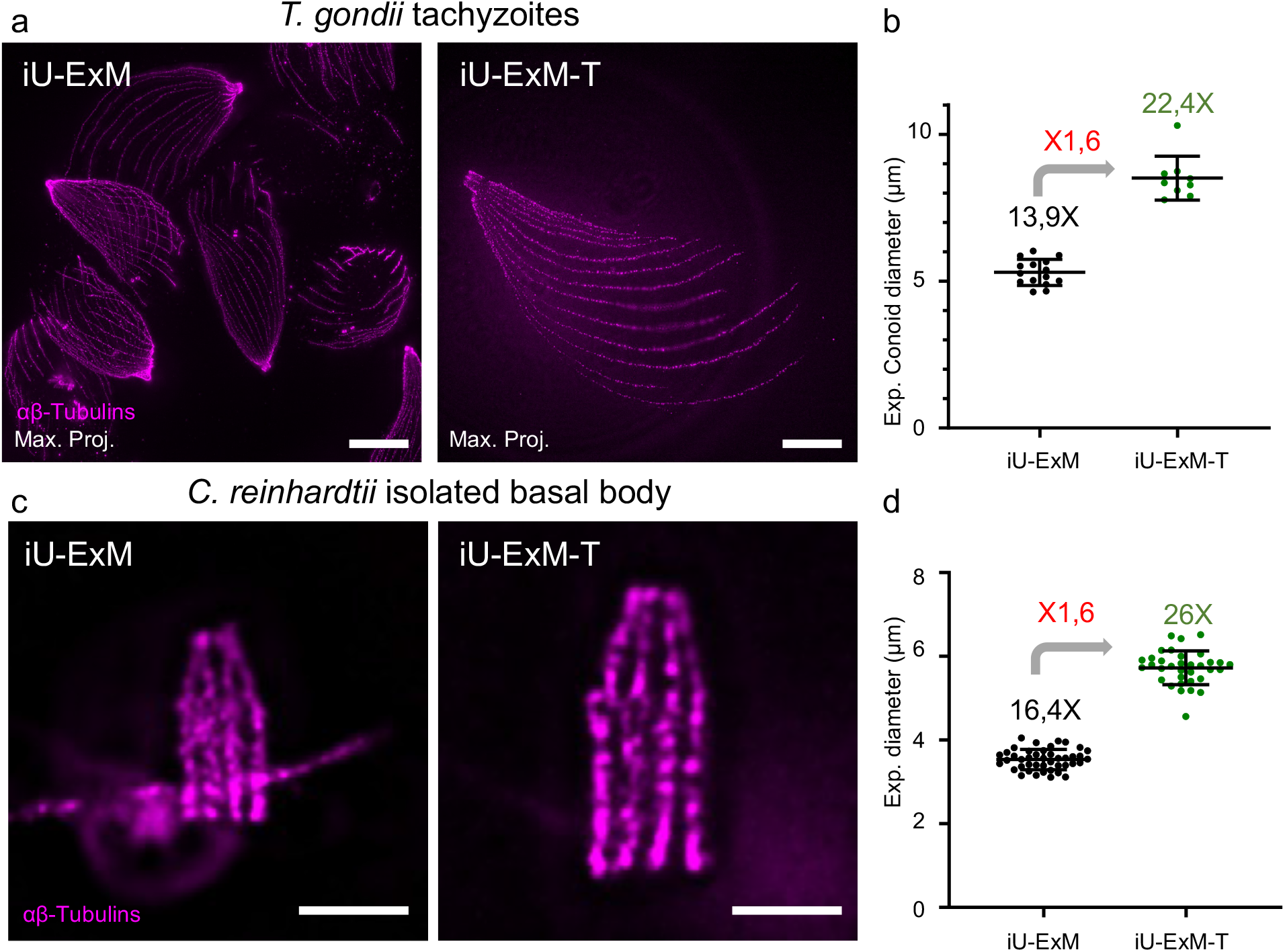
Coupling TREx to iU-ExM. (**a)** Widefield full field of view of *T. gondii* tachyzoites stained for tubulin (magenta) expanded with either iU-ExM (left) or iU-ExM-T (right). Scales bar: 30 μm non-corrected. (**b)** Quantification of the diameter of the conoid, highlighting the 1.6-fold improvement on the expansion factor. iU-ExM: N= 15 conoids (average +/- standard error = 5.3 +/- 0.4 μm), iU-ExM-T: N= 9 (M= 8.5 +/- 0.7 μm). Data from one experiment. (**c)** Representative widefield image of *C. reinhardtii* isolated basal bodies stained for tubulin (magenta) expanded using iU-ExM (right) or iU-ExM-T (left). Scale bars: 5 μm non-corrected. (**d)** Quantifications of the centriole diameter in the proximal region, further illustrating the 1.6-fold improvement of the expansion factor. iU-ExM: N= 43 centrioles (average +/- standard error = 3.5 +/- 0.2 μm), iU-ExM-T: N= 34 (average +/- standard error = 5.7 +/- 0.4 μm) centrioles from one experiment.

**Supplementary Figure 7.**
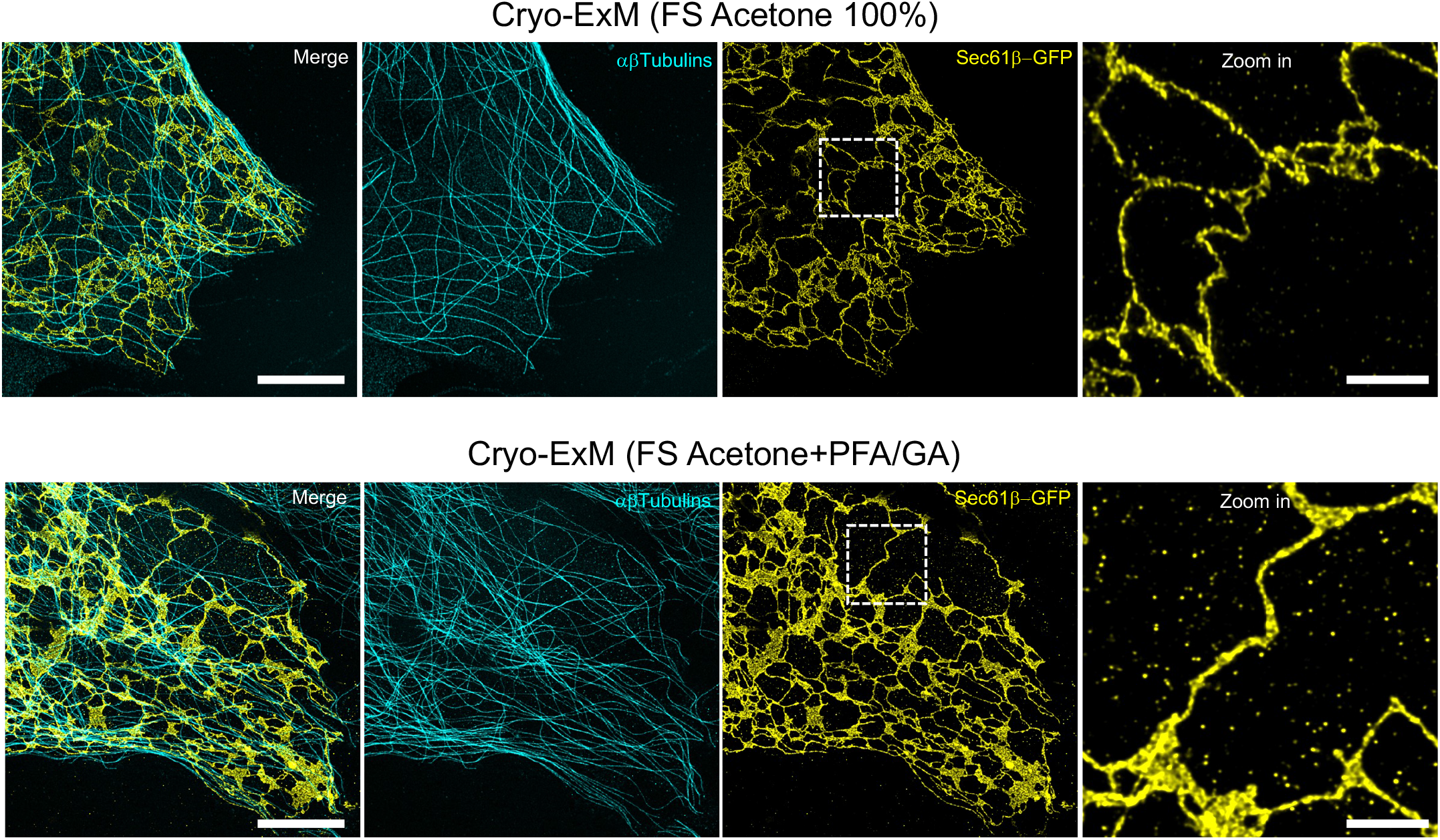
Endoplasmic reticulum membrane preservation in Cryo-ExM in presence of 0.02% glutaraldehyde. **(a)** Confocal image of an expanded Sec61β-GFP transfected U2OS cell after cryo-fixation and freeze substitution (FS) in acetone 100%. Cells were stained for anti-GFP (yellow) and anti-tubulin (cyan) antibodies labelling the endoplasmic reticulum (Sec61β-GFP) and the microtubules respectively. White dashed square indicates the inset (zoom in) showing a slightly collapsed endoplasmic reticulum in 100% acetone FS conditions. Scale bars: 5 mm and 1 mm (inset). **(b)** Confocal image of an expanded U2OS after cryo-fixation and freeze substitution (FS) in acetone supplemented with PFA 0.5% and GA 0.02%. Cells were stained for anti-GFP (yellow) and anti-tubulins (cyan) antibodies. White dashed square indicates the inset (zoom in) showing a non-collapsed endoplasmic reticulum in PFA/GA FS conditions. Scale bars: 5 mm and 1 mm (inset).

**Supplementary Figure 8.**
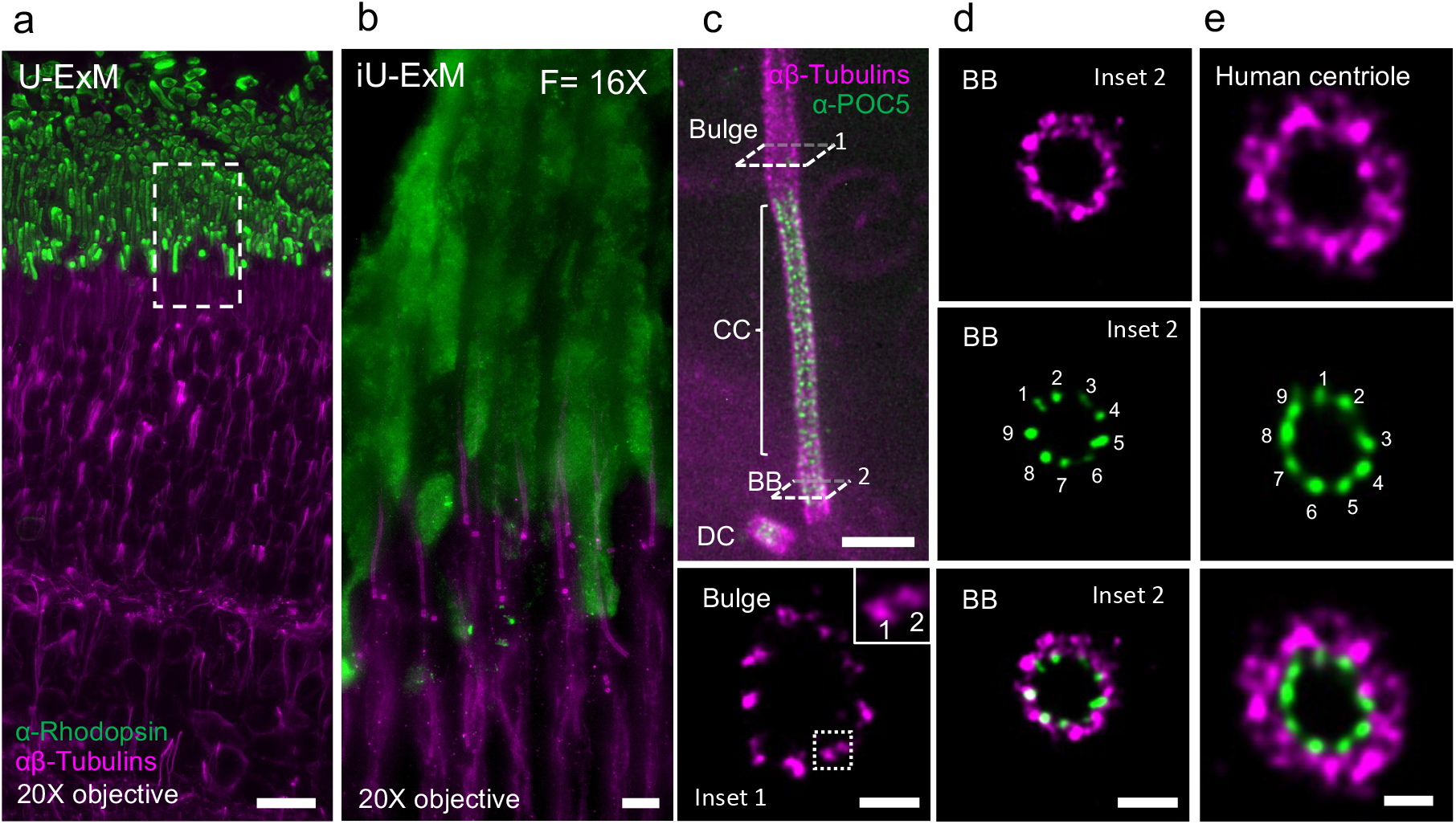
iU-ExM on retinal mouse tissue. (a). U-ExM widefield image of an expanded mouse retinal tissue stained for anti-tubulins (magenta) and anti-rhodopsin (outer segment, green) using a 20x air objective. Scale bar: 10 μm. **(b)** iU-ExM widefield image of an expanded mouse retinal tissue stained for tubulin (magenta) and rhodopsin (outer segment, green) using a 20x air objective highlighting the gain in expansion and resolution. Scale bar: 1 μm. (**c)** Top: widefield image of a iU-ExM expanded photoreceptor cell stained for tubulin (magenta) and POC5 (green) highlighting the connecting cilium (CC) region. BB: Basal Body, CC: connecting cilium, DC: Daughter centriole. Scale bar: 500 nm. Inset 1: confocal top view of the bulge region of a iU-ExM expanded photoreceptor. Note that individual microtubule blades from the microtubule doublets are now distinguishable. Scale bar: 200 nm. (**d)** Confocal top view of a basal body underlying the CC of a iU-ExM expanded photoreceptor cell stained for tubulin (magenta) and POC5 (green), unveiling the 9-fold organization of POC5. Note that the tubulin signal at the level of BB is less clear than that at the CC (Fig. 3i). Scale bar: 200 nm. (**e)** Top view confocal picture of a human centriole from U2OS cells labelled with tubulin and POC5 showing the conservation of the 9-fold pattern in human centrioles. Note also that similarly to the BB, the tubulin signal is not nicely resolved at centrioles in human cells. Scale bar: 200 nm.

**Supplementary Figure 9.**
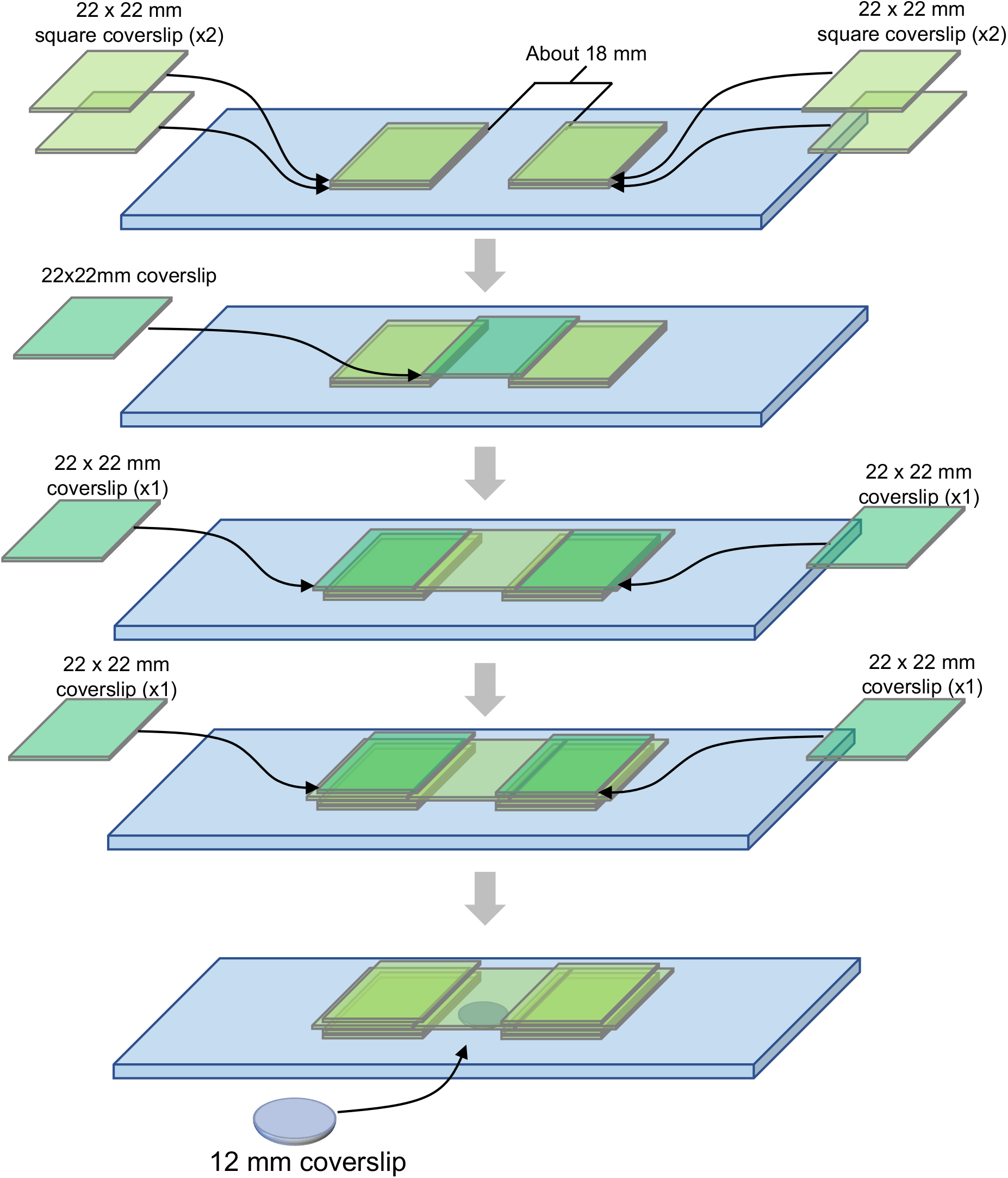
Scheme of the gelation chamber.

**Movie 1**. Widefield video of top view NPCs from iU-ExM expanded purified NUP96-GFP (red hot) nuclei. The sample was stained with α-GFP and α-NUP96 antibodies and imaged at 100X with a 1.4NA objective. Scale bar: 1 μm.

**Movie 2**. Widefield video of NPC in side view from iU-ExM expanded purified NUP96-GFP (red hot) nuclei. The sample was stained with α-GFP and α-NUP96 antibodies and imaged at 100X with a 1.4NA objective. Scale bar: 1 μm.

## Reproducibility

All experiments were performed at least 3 times with some exceptions: iU-ExM-T on *T. gondii* tachyzoites: N= 1. DCX staining on *T. gondii* tachyzoites after 73°C denaturation: N= 1. *T. gondii* tachyzoites after 73°C denaturation labelled with tubulin: N= 2. iU-ExM on NUP96-eGFP U2OS pre-extracted cells labelled with α-NUP96 antibody only: N= 1.

## Data availability statement

The data that support the findings of this study are available from the corresponding authors upon request. Request regarding *T. gondii* strains should be adressed to DSF.

## ONLINE METHODS

## Reagents

| Product | Supplier | Reference |
| --- | --- | --- |
| N,N'-(1,2-Dihydroxyethylen)-bisacrylamid (DHEBA) | Merck – Sigma Aldrich | 294381 |
| Bisacrylamide (BIS) | Merck – Sigma Aldrich | M1533 |
| Acrylamide 40% w/w | Merck – Sigma Aldrich | A4058 |
| Formaldehyde 35-38% | Merck – Sigma Aldrich | F8775 |
| Sodium acrylate | AK Scientific | R624 |
| Sodium Hydroxyde | Merck – Sigma Aldrich | 206060010 |
| Amonium persulfate (APS) | Thermo Fisher | 17874 |
| Tetramethylethylenediamine (TEMED) | Thermo Fisher | 17919 |
| DMEM supplemented with glutamax | Thermo Fisher | 61965-026 |
| Trypsin | Thermo Fisher | 25300-054 |
| Fetal Bovine Serum | Thermo Fisher | 10270 |
| Penicillin-Streptovidine | Merck – Sigma Aldrich | R8758 |
| Poly-D-Lysine | Merck – Sigma Aldrich | A38904-01 |
| Sodium Dodecyl Sulfate | Pan Reac Applichem | A7219 |
| Tris-Base | Roth | 2449.3 |
| NP-40 | Merck – Sigma Aldrich | I8896 |
| Tween 20 | Roth | 9127-2 |
| Nuclease Free water | Invitrogen | AM9937 |
| Bovine Serum Albumine (BSA) | Merck – Sigma Aldrich | 10735086001 |
| Twinsil Picodent | Picodent | Twinsil |
| Glutaraldehyde | Merck – Sigma Aldrich | G5882 |
| Paraformaldehyde 16% | Electron Microscopy Science | 15710 |
| Antibody/Dye | Supplier-Reference | Dilution/Concentration |
| $\alpha$ Tubulin monobody | AA344 | 1:100 (iU-ExM), 1:250 (U-ExM) |
| $\beta$ Tubulin monobody | AA345 | 1:100 (iU-ExM), 1:250 (U-ExM) |
| HsPOC5 | Bethyl – A303-341A | 1:200 (iU-ExM & U-ExM) |
| GFP | Torrey Pine-TP401 | 1:200 (iU-ExM), 1:250 (U-ExM) |
| NUP98-96 | ProteinTech-12329-1-AP | 1:200 (iU-ExM) |
| NUP205 | ProteinTech-24439-1-AP | 1:200 (iU-ExM) |
| Rat Anti-HA | ROCHE – 11 867 423 001 | 1:250 (iU-ExM) |
| Rhodopsin | Thermoscientific-MA5-11741 | 1:250 (iU-ExM) |
| WGA CF568 | Biotium-29077-1 | 20 µg/mL (iU-ExM) |
| NHS-Ester | Atto-Tec-AD594-31 | 20 µg/mL(iU-ExM) |
| Rat 488 | Invitrogen – A21208 | 1:250 (iU-ExM), 1:400 (U-ExM) |
| Rabbit 488 | Invitrogen-A11008 | 1:250 (iU-ExM), 1:400 (U-ExM) |
| Mouse 488 | Invitrogen-A11029 | 1:250 (iU-ExM), 1:400 (U-ExM) |
| Mouse 568 | Invitrogen-A11004 | 1:250 (iU-ExM), 1:400 (U-ExM) |
| Guinea Pig 568 | Invitrogen-A11075 | 1:250 (iU-ExM), 1:400 (U-ExM) |

### Cell lines and strains used in this study

#### Chlamydomonas reinhardtii

To unsure an efficient expansion, we used a cell wall free *C. reinhardtii* CW15^-^ strain. The liquid culture (TAP buffer ^31^) is inoculated from solid agarose culture of the algae and grown for 3 days at RT under light exposure and slow shaking. The cells were sedimented on 12 mm Poly-D-Lysine coated coverslips for 10 min, then the excess was removed and the cells fixed with either cold methanol (see cell fixations) or no fixation.

#### Toxoplasma gondii

*Toxoplasma gondii* tachyzoites were amplified in human foreskin fibroblasts (HFFs, ATCC) in Dulbecco’s Modified Eagle’s Medium (DMEM, Gibco) supplemented with 5% of Fetal Bovine Serum (FCS, Gibco), 2 mM glutamine and 25 µg/ml gentamicin (Gibco). Tachyzoites were transfected by electroporation 65. To target any gene of interest, 40 µg of specific gRNA was transfected alongside a PCR product flanked by homology regions (gRNA sequencies: DCX: GTGGGGAGCGTGTCACTCAT, SAS6-L: ACTTATGTACGAGTGCACGG.

Parasites carrying an HXGPRT cassette^32^ were selected with 25 mg/ml of mycophenolic acid and 50 mg/ml of xanthine. Parasites carrying a DHFR cassette (doi: https://journals.asm.org/doi/10.1128/mBio.01114-14) were selected with 1 µg/ml of pyrimethamine. For immuno-staining as well as for U-ExM and iU-ExM, parasites were sedimented for 10 min on pre-coated Poly-L-Lysine coverslips before starting the protocol described in this study.

### Human cell lines

*Homo sapiens* bone osteosarcoma U2OS ATCC-HTB-96 and NUP96-GFP U2OS cell line (from Jonas Ries lab ^13^) were grown in Dulbecco’s modified Eagle’s medium and GlutaMAX, supplemented with 10% fetal calf serum and penicillin and streptomycin (100 μg/ml) at 37 °C in a humidified 5% CO_2_ incubator. U2OS expressing GFP-sec61β (133-291) were transiently transfected with JetPRIME following the manufacturer’s instructions. After 24h of expression, cells were cryo-fixed as described in ^25^. All cell cultures were regularly tested for mycoplasma contaminations.

### Isolated *Chlamydomonas reinhardtii* Basal Bodies/Centrioles

The purified basal bodies were prepared and spun on coverslip for expansion as previously described ^31^. Briefly, deflagellated CW15-*Chlamydomonas* cells were lysed 1h at 4°C in presence of 1 mM HEPES (pH 7), 0.5 mM MgCl2, 1% NP-40, and 5000 units of DNase. After centrifugation at 600g for 10 min at 4°C to remove the cell debris, basal bodies were further purified and concentrated, first using a centrifugation at 10,000g with a 60% sucrose cushion and second a centrifugation at 68,320g on a 40% to 70% sucrose gradient. Isolated basal bodies were collected at the 70% sucrose interface.

### Cell fixations

According to the type of structure that we investigated, different fixations were used:

### Glutaraldehyde-Formaldehyde fixation

Immediately after removing the plate from the incubator, the medium was removed from the well and 4 mL of fixing solution (0.1% Glutaraldehyde (GA); 3% formaldehyde (FA) in PBS 1x) was poured for 15 min at RT or at 37°C (for the mitochondria) in the well without any rinsing to insure proper fixation. Next, cells were rinsed 3 times with PBS and expanded shortly after.

### Cold Methanol fixation

The coverslips with cells were plunged into cold methanol (−20°C) and incubated in the freezer (−20°C) for 7 minutes.

### Cryo-fixation

U2OS cells were cryo-fixed, and freeze substituted as previously described^25^, with some minor modifications. Briefly, the coverslips containing the sample were held halfway with a thin tweezer (Dumont 5, Sigma F6521-1EA). After blotting the remaining medium, the coverslips were plunged with a homemade plunge freezer an ethane/propane mix cooled with liquid nitrogen. Coverslips were transferred into a 5-ml Eppendorf tube containing 1.5 ml of liquid nitrogen-chilled acetone supplemented with paraformaldehyde-Glutaraldehyde (PFA-GA) at 0.5-0.02% repectively. Tubes were placed on dry ice with a 45° angle and agitated overnight to allow the temperature to rise to −80 °C and further incubated without dry ice for 1h until the temperature reached ∼0 °C. Samples were then rehydrated in successive ethanol: water solutions supplemented with PFA-GA (0.5-0.02%), as follows: ethanol 100%, ethanol 100%, ethanol 95%, ethanol 95%, ethanol 70%, ethanol 50%, ethanol 25% and PBS. Note that coverslips were incubated in the ethanol 100% solutions for 5 min and 3 min for the following. Cells were directly processed for expansion.

## iU-ExM protocol

### Gelation chamber

On a glass microscope slide, two stacks of two 22 × 22 mm coverslips (no. 1.5) were glued to the slide with enough space for a 12 mm round coverslip in between. Next, one 22 × 22 mm coverslips was added as a lid and secured with 2 coverslips on the side and two on top to create a rack where the lid coverslip can slide in. In this configuration, the lid coverslip cannot move, and the gel thickness would be approximatively 170 μm (**Supplementary Figure 9**).

### First Expansion

#### Fixation

Dependent on the sample or imaged organelle. See cell fixations.

#### Anchoring

The sample was incubated in anchoring solution (2% AA; 1.4% FA in 1x PBS) for 3h at 37°C.

#### Gelation

After anchoring, the coverslip was slightly dried with Kimwipes and sealed in the gelation chamber (**Supplementary Fig. 9**), then put on a humid chamber on ice. Using a gelation chamber is important to insure a controlled and homogeneous height of the gel (around 170 μm). Next, a monomer solution (MS) (10% AA, 19% Sodium Acrylate (SA), 0.1% DHEBA, 0.25% tetramethylethylenediamine (TEMED)/Amonium Persulfate (APS)) was added to fill the space between the coverslip and the lid of the gelation chamber so that it covers entirely the 12 mm coverslip. After 15 min on ice, the humid chamber was placed at 37°C for 45 min to complete the gelation.

#### Denaturation

After gelation, the coverslip with the gel was carefully removed from the imaging chamber and dipped in 2 mL of denaturation buffer (200 mM Sodium Dodecyl Sulfate (SDS); 200 mM NaCl; 50 mM Tris-BASE; pH=6.8) in a 6 well plate under shaking until the gel detach from the coverslip. Next, the gel was transferred in a 1.5 mL Eppendorf tube with 1mL of fresh denaturation buffer and incubated for 1h30 at 85°C. The temperature was carefully watched with an external thermometer. A few degrees below can cause expansion anisotropy while few degrees higher can cause gel disintegration.

#### 1^st^ Expansion step

after denaturation at 85°C, the gel was dipped into ddH2O in a 12 cm petri dish. The water was changed every 20-30 min until the expansion of the gel plateaus. We observed that the expansion factor is around 5 -6X, depending on the crosslinker purity, pH of the denaturation buffer, temperature and/or time of denaturation. Note that the DHEBA crosslinker is sensible to pH and temperature, and at 85°C some crosslinkers are cleaved.

#### Intermediate antibody staining

he immunolabelling is performed after the 1^st^ expansion step (see staining procedures).

### 2^nd^ expansion

#### Neutral gel embedding

The first expanded gel was cut in 1 cm^2^ piece and placed in a 6 well plate on ice. 3 washes with activated neutral gel (10% AA; 0.05% DHEBA; 0.1% APS/TEMED in ddH2O) for 10 min under shaking were done. Due to the APS salt content, the gel is expected to shrink about 1.5x. The gel was then put on a microscope slide, and the excess of monomer solution was gently removed with Kimwipes and the gel covered by a 22 × 22 mm coverslip and lastly incubated in a humid chamber for 1h at 37°C.

#### 2^nd^ anchoring

Note that while this step can be avoided in the Pan-ExM original protocol ^9^, it is required for iU-ExM to retain the antibody staining in the 2^nd^ gel. The gel embedded in the neutral gel was incubated in anchoring solution (1.4% FA/2% AA) for 3-5h under shaking at 37°C. The gel was then washed in PBS 1x for 30 minutes.

#### 2^nd^ monomer solution embedding

In a 6 well plate, the gel was washed 3 times for 10 min under shaking and on ice with the 2^nd^ expansion monomer solution (10% AA, 19% SA, 0.1% BIS, 0.1% TEMED/APS) for a ± 16x expansion (iU-ExM) or the TREx^23^ monomer solution (14.5% AA; 10.5% SA; 0.01%BIS; 0.1% APS/TEMED in ddH_2_O) giving rise to a 24-26x linear expansion (iU-ExM-T). Next, the excess monomer solution was gently removed with Kimwipes and the gel covered by a 22 × 22 mm coverslip and incubated in a humid chamber for 1h at 37°C. After final polymerization, the gel was incubated in 200 mM NaOH solution for 1h under shaking at RT followed by washes of ± 20 min with PBS 1x until the pH drops to 7. The gel was next dipped in ddH_2_O and the water was changed until the expansion of the gel plateaus.

#### Adaptation for mouse retina tissue expansion

The 1^st^ expansion of the sample was processed as described in ^27^. Briefly, in a 35 mm Mattek dish with 10 mm micro well (P35G-1.5-10-C), the retina was first embedded in anchoring solution (1.4% FA/2% AA) overnight at 37°C. After that, the tissue was incubated for 45 min in a monomer solution (19% SA; 10% AA; 0,2% DHEBA in 1x PBS) without APS/TEMED to insure a proper diffusion of the monomer solution in the tissue. Note that the DHEBA concentration is doubled to strengthen the gel to insure proper expansion. Indeed, as the denaturation is longer, more crosslinkers will be cleaved and the gel will be too brittle to be processed. Then fresh activated MS was added, and a 22 * 22 mm coverslip was placed on top of the Matek well for 45 min on ice followed by 1h at 37°C in a humid chamber. Denaturation buffer was next added on the polymerized gel until the gel pops out of the well. Next, the gel was placed on a 1.5 mL Eppendorf with fresh 1 mL of denaturation buffer and incubated at 85°C for 2h. The gel was then expanded in ddH_2_O and sliced as previously described^27^. The 2^nd^ expansion was performed on the slices as described for the regular iU-ExM protocol.

## Pan-ExM Protocol

The Pan-ExM protocol was done has described ^33^. Briefly, to analyse centrioles, cells were incubated without fixation in 0.7% formaldehyde + 1% acrylamide (w/v) in 1× PBS for 6h at 37°C. After washing the cells in PBS, in a gelation chamber (**Supplementary fig. 9**) cells were incubated in the monomer solution containing (19% (w/v) sodium acrylate (SA) + 10% acrylamide (AA) (w/v) + 0.1% (w/v) DHEBA (N,N′-(1,2-dihydroxyethylene) bisacrylamide) + 0.25% (v/v) TEMED (N,N,N′,N′-tetramethylethylenediamine) + 0.25% (w/v) Ammonium persulfate (APS) in PBS and incubated for 1h at 37°C in a humid chamber to reach complete polymerization. Then, the gel was dipped in 2 mL denaturation buffer under shaking until the gel detaches from the coverslip. The gel was next transferred in a 1.5 mL Eppendorf tube with 1 mL of fresh denaturation buffer and incubated for 1h at 73°C. Then, the gel was expanded in ddH_2_O with at least 3 washes, until the expansion of the gel plateaus. The gel should expand 4-4.5X according to the DHEBA purity. Then, the gel was cut in a 1 cm^2^ piece and embedded in a neutral gel (10% AA; 0.05% DHEBA; 0.05% APS/TEMED in ddH2O). Embedded gels were incubated in a third monomer solution containing (19% (w/v) SA + 10% AA (w/v) + 0.1% (w/v) BIS + 0.05% (v/v) TEMED + 0.05% (w/v) APS in PBS). After polymerization, the first gel containing DHEBA crosslinkers was dissolved by incubating it in 0.2M NaOH for 1h. After several washes, the gels were subjected to immunostaining (see immunostaining below), and placed in distilled water for the final expansion procedure.

## Ultrastructure expansion microscopy (U-ExM)

Expansion of unfixed and cryo-fixed cells was performed as previously described ^6,25^. Cells were incubated for 3h in anchoring solution (2% AA, 1.4% FA in 1x PBS) at 37 °C before gelation in U-ExM monomer solution (10% AA, 19% SA, 0.1% BIS in 1x PBS) containing 0.5% TEMED and APS. Next, cells were incubated for 5 min on ice followed by 1 h at 37 °C and incubated for 1h30 at 95 °C in denaturation buffer. Gels were washed from the denaturation buffer twice in ddH_2_O.

## Staining procedures

### Immunostainings

#### iU-ExM intermediate staining/U-ExM gels

the first gels were shrunk in 1x PBS and stained for 3h at 37°C in 1x PBS-BSA 2% for both primary and secondary antibodies, followed each by 3 washes for 15 min with 1x PBS-Tween 0.1%. For NPCs, the primary antibody staining was done overnight at 4°C and secondary for 6h at 37°C, followed each by 3 washes for 15 min with 1x PBS-Tween 0.1% (see table for concentrations and antibody reference). The gel was next re-expanded in ddH_2_O.

#### iU-ExM/Pan-ExM post staining

The gel was first shrunk in 1x PBS. The primary antibodies were diluted in 1x PBS – BSA 2% (see table for the dilutions). The gel was next incubated with the antibodies for at least 12h at 37°C under shaking. The secondary antibodies were diluted in 1x PBS-BSA 2% and incubated with the gel for 6h-12h minimum at 37°C. Lastly, the gel was washed 3 times for 30 min minimum with 1x PBS-Tween 0.1% before imaging.

### Other labelling

#### WGA staining

For iU-ExM, the WGA staining is done after the antibody labelling on the 1^st^ gel. The gels are shrunk in 1x PBS and incubated under shaking at 37°C for 1h30 min with 10 μg/mL of WGA CF-568 in 1x PBS followed by 3 washes for 15 min with 1x PBS, Tween 0.1%.

#### NHS-Ester staining

The gels (from either U-ExM, iU-ExM or Pan-ExM protocols) were incubated 1h30 at RT under shaking with NHS-Ester ATTO 594 diluted at 20 μg/mL in 1x PBS. The gels were next washed 3 times with PBS-Tween 0.1% for at least 30 min under shaking. Note that the NHS-Ester staining post-expansion can lead to some unspecific signal by labelling the antibodies used for the intermediate staining procedure. For U-ExM/first expanded iU-ExM/Pan ExM gels, the NHS-Ester staining is done after immuno-labelling because NHS-Ester binding might cover the epitopes.

#### DAPI staining

The gel was shrunk in 1x PBS then stained with DAPI at 1 μg/mL in 1x PBS for 15 min under shaking at RT followed by 3 washes with PBS-Tween 0.1%. Note that for iU-ExM gels, the DAPI staining shows better intensity when labelled with intermediate staining procedure.

## Gel mounting

For all stage of expansion (1^st^ or 2^nd^), gels were cut with a razor blade into squares to fit in a 36 mm metallic imaging chamber. The excess water was carefully removed with kimwipes, being careful not to dry it too much to prevent shrinking and the gel as placed on a Poly-D-Lysine coated 24 mm coverslips to prevent drifting. The 2^nd^ expanded gels were prone to drifting even on Poly-D-Lysine coated coverslips. If drifting too much, the gel was stabilized by being embedded in a Twinsil Picodent in the chamber.

## Image acquisition & analysis

Confocal and widefield images were acquired using a Leica SP8 and a Leica Thunder using a 63x /1.4 NA or 100x/1.3 NA oil immersion objectives. Microscopes parameters are controlled using the Suite X software (LAS X; Leica Microsystems). The images were deconvolved with either LVCC for large image or SVCC for small ROI with widefield microscope and with Lightning deconvolution for confocal images. All the deconvolution algorithm are from Leica Microsystems. Images were processed and quantifications were done with ImageJ (FIJI)^28^. Before mounting, images are smoothed with Fiji.

## Gel conservation

Either stained or non-stained expanded gels can be stored for at least a year in 50% glycerol at-20°C. Adding glycerol prevents the gel from freezing. For storage, the expanded gel should be washed at minimum 3 times for 1h with 50% glycerol, until the glycerol completely diffuses in the gel. Due to the difference of diffraction index between water and glycerol, if the diffusion of the glycerol is not complete the gel will remain visible in the glycerol solution and will froze. To defrost the gels, place them in 1x PBS where they will shrink and expulse all the glycerol. Next, the gels should be washed 2 times for 30 min minimum with PBS1x and then expanded in ddH2O at least once before staining. Insufficient wash would cause weak and noisy antibody labelling for unstained gel and optic aberrations for stained gel due to the altered refractive index due to the presence of glycerol.

## Quantifications

### Nuclei area measurement

When the intensity of the DAPI allows it (mostly for 1^st^ expanded gels and non-expanded cells), the images were automatically binarized and the particles were selected with the particle tool of imageJ with a home-made imageJ plugin. Then, the selected areas were controlled by hand: deformed, overlapping, part out of the picture, were excluded. For iterative gels, when the DAPI staining did not allow automatic segmentation of the nuclei, area was manually measured on ImageJ ^34^. As the nucleus can be distinguishable with NHS-Ester or Sec61β staining, the nuclei area could also be manually measured with those staining (at least 10 nuclei).

### Nuclear Pore Complex analysis

The coverslips with cells were prepared as follow: first the cells were pre-fixed with 2.4% FA in PBS for 30 seconds and permeabilized for 3 min with 0.4% Triton X-100 in 1x PBS and washed 2 times with PBS for 5 minutes. Then, cells were fixed with 2.4% FA in PBS for 20 minutes at RT and washed 2 times for 5 min with 1x PBS. Finally, a second permeabilization with 0.2% Triton X100 in 1x PBS was performed for 10 min and followed by 2 washes of 1x PBS for 5 min. The coverslips were then stored in 1x PBS before expansion.

#### Nuclei purification

To separate the nuclei from the cytoplasm, we followed a previously published protocol^29^. Briefly, the cells were harvested from a T75 flask with trypsin and centrifuged at 1000 rcf for 5 min. The pellet was resuspended in 5 mL of complete DMEM and aliquoted with 1mL in 1.5 mL Eppendorf tubes. Cells were pelleted with 1000 rcf for 5 min and resuspended in hypotonic buffer (20 mM Tris-HCl pH 7.4, 10 mM KCl, 2 mM MgCl_2_, 1 mM EGTA, 0.5 mM DTT, 0.5 mM PMSF) for 3 min on ice. Then, NP-40 was added to a final concentration of 0.3% for 3 min on ice to lyse the cells with vortexing. The suspension was next centrifuged at 1000 rcf for 5 min and the supernatant (cytoplasmic material) removed. The pellet (nuclei) was resuspended in isotonic buffer (20 mM Tris-HCl pH 7.4, 150 mM KCl, 2 mM MgCl_2_, 1 mM EGTA, 0.5 mM DTT, 0.5 mM PMSF) supplemented with NP-40 to reach 0.3% final concentration for 3 min on ice with vortexing to further purify the nuclei. Nuclei were then pelleted at 1000 rcf for 5 min and resuspended in 1x PBS. Finally, purified nuclei were pelleted on 12 mm Poly-D-lysine coverslips at 1000 rcf for 15 min, by adding one 12 mm coverslip on a well with 500 μL of nuclei preparation. The nuclei were then fixed with 2.4% FA in 1x PBS for 20 min at RT and the coverslips were either expanded or mounted for regular immunofluorescence.

#### Angle measurement

To measure the angle between two consecutive NUP96 dots, only the most circular and in a good focal plane NPCs were selected (titled NPCs and NUP96 dots closer than 20° were excluded). In the case of a partial staining (less than 8 dots), only consecutives dots angle were measured.

#### Labelling efficiency

To count the number of NUP96 spots per NPC, only NPC in a good focal plan and displaying a circular WGA signal was manually selected. Then, distinct dots were manually counted.

## Expansion factor measurement

For double expanded gels it is not possible to have the expansion factor by measuring the gel because of all the intermediary steps that add too much variability. Thus, it is more accurate to rely on a biological indicator. We assumed that the potential linkage error due to antibody labelling is neglectable regarding the expanded size of the structure. However, it is important to state that the corrected measurements are subjected to the ground truth value used to measure the expansion factor.

To obtain the expansion factor: for membranous structures (mitochondria, endoplasmic reticulum), the expended cross section of the nuclei was divided by the non-expanded average cross-section. To obtain the NPC expansion factor, the diameter of the NUP96 signal was measured and divided by 107 nm as previously published ^13^ to obtain the expansion factor that will be further used for all the other measurements. Between a NUP96 and NUP205 staining of a same experiment, we controlled that the expanded WGA diameter was identical before extending the calculated expansion factor to the NUP205 measurements. For *Toxoplasma gondii*, the expanded diameter of the conoid at the apical part was divided by 380 nm ^19^. For *C. reinhardtii* centrioles, the proximal diameter is divided by 225 nm ^30^. For retina expansion, the expanded diameter of the connecting cilium was divided by 200 nm, for 50% maximal intensity ^27^.

## Statistical analysis

The normality distribution of every data set was assessed with Shapiro-Wilk test. If normality passed, ANOVA or student test was runed. If not, non-parametric statistical analysis was done on Graphepad Prism assuming equal SD. N indicates independent biological replicates from distinct samples. Data are all represented as scatter dot plots with center line as mean. The graphs with error bars indicate 1 SD (+/-) and the significance level is denoted as usual (*p<0.05, **p<0.01, ***p<0.001). All the statistical analyses were performed using Prism7 (Graphpad version 7.0a, April 2, 2016).

## Notes

### Competing Interest Statement

The authors have declared no competing interest.

